# Efficient CRISPR/Cas-mediated Targeted Mutagenesis in Spring and Winter Wheat Varieties

**DOI:** 10.1101/2021.05.09.443285

**Authors:** Florian Hahn, Laura Sanjurjo Loures, Caroline A. Sparks, Kostya Kanyuka, Vladimir Nekrasov

**Affiliations:** Plant Sciences Department, Rothamsted Research, Harpenden AL5 2JQ, United Kingdom; Department of Biointeractions and Crop Protection, Rothamsted Research, Harpenden AL5 2JQ, United Kingdom

**Author notes:** Department of Plant Sciences, University of Oxford, Oxford OX1 3RB, United Kingdom (Florian Hahn).

**Keywords:** CRISPR, Cas9, plant, genome editing, BAK1, eIF4E, wheat

## Abstract

The CRISPR/Cas technology has recently become a molecular tool of choice for gene function studies in plants as well as crop improvement. Wheat is a globally important staple crop with a well annotated genome and there is plenty of scope for improving its agriculturally important traits using genome editing technologies, such as CRISPR/Cas. As part of this study we targeted three different genes in hexaploid wheat *Triticum aestivum*: *TaBAK1-2* in the spring cultivar Cadenza as well as *Ta-eIF4E* and *Ta-eIF(iso)4E* in winter cultivars Cezanne, Goncourt and Prevert. The primary transgenic lines carrying CRISPR/Cas-induced indels were successfully generated for all targeted genes. As winter wheat varieties are generally less amenable to genetic transformation, the successful experimental methodology for transformation and genome editing in winter wheat presented in this study will be of interest to the research community working with this crop.

## Introduction

Common wheat (*Triticum aestivum*) is one of the most important staple food crops in the world. The challenges that global agriculture currently faces, such as growth of the world’s population and climate change, dictate demand for technologies with a potential to accelerate crop breeding (Reynolds et al., 2021). During the last decade, genome editing emerged as a powerful new plant breeding technique (NBT) (Holme et al., 2019) that enables targeted changes in crop genomes.

CRISPR/Cas is by far the most common plant genome editing technology nowadays due to its versatility and ease of use (Kumar et al., 2021). Bread wheat is an allohexaploid, therefore it is important to have an efficient CRISPR/Cas setup as in the majority of cases, for each particular gene, one needs to target six copies i.e. two per each of the three subgenomes (A, B and D).

As part of this study, we used CRISPR/Cas in a reverse genetics approach to target the *TaBAK1-2* gene, a homologue of the Arabidopsis *BAK1* gene encoding the BRI1-ASSOCIATED RECEPTOR KINASE 1 (BAK1) – an important regulator of plant immunity and development (Yasuda et al., 2017; Nolan et al., 2020), in the spring cultivar Cadenza. Here we successfully knocked out all three *TaBAK1-2* homoeologues in primary transgenic lines and demonstrated transmission of the CRISPR/Cas-induced mutant alleles to the next generation (T1). We anticipate the resultant homozygous mutant lines will facilitate studies on the involvement of BAK1 in immune responses in wheat.

In the second part of the study, we tested the potential of the CRISPR/Cas system in wheat for generating resistance to bymoviruses in the family Potyviridae, some of which are serious pathogens of crops. For instance, *Wheat spindle streak mosaic virus* (WSSMV) can pose a serious threat to wheat production in Europe and North America, while *Wheat yellow mosaic virus* (WYMV) – in East Asia (Jiang et al., 2020). Here, we targeted *Ta-eIF4E* and *Ta-eIF(iso)4E* genes encoding highly conserved translation initiation factors eIF4E and eIF(iso)4E, respectively, which serve as susceptibility (S) factors required for plant viruses from the *Potyviridae* family to complete their life cycle (Revers and García, 2015). An analogous genome editing based strategy has already been successfully used in Arabidopsis, cucumber and cassava (Schmitt-Keichinger, 2019). In addition, in barley, the conventional breeding strategies for generating resistance to bymoviruses *Barley mild mosaic virus* (BaMMV) and *Barley yellow mosaic virus* (BaYMV) are based on introducing recessive mutant alleles of the *eIF4E* gene (Kanyuka et al., 2005; Jiang et al., 2020). In this study, we generated genome edited wheat lines carrying indels in all three homoeologues of either *TaeIF4E* or *TaeIF(iso)4E*. These lines will be assessed for enhanced resistance to WSSMV in the follow-up study.

## Material and Methods

### Target sites

The sgRNA target sites were chosen using the CRISPOR online tool (Concordet and Haeussler, 2018) or Geneious software. The target genes were sequenced in all wheat varieties used for transformation to ensure the presence of the chosen sgRNA target sites (Table 1) in each variety and in each subgenome.

**Table 1.**
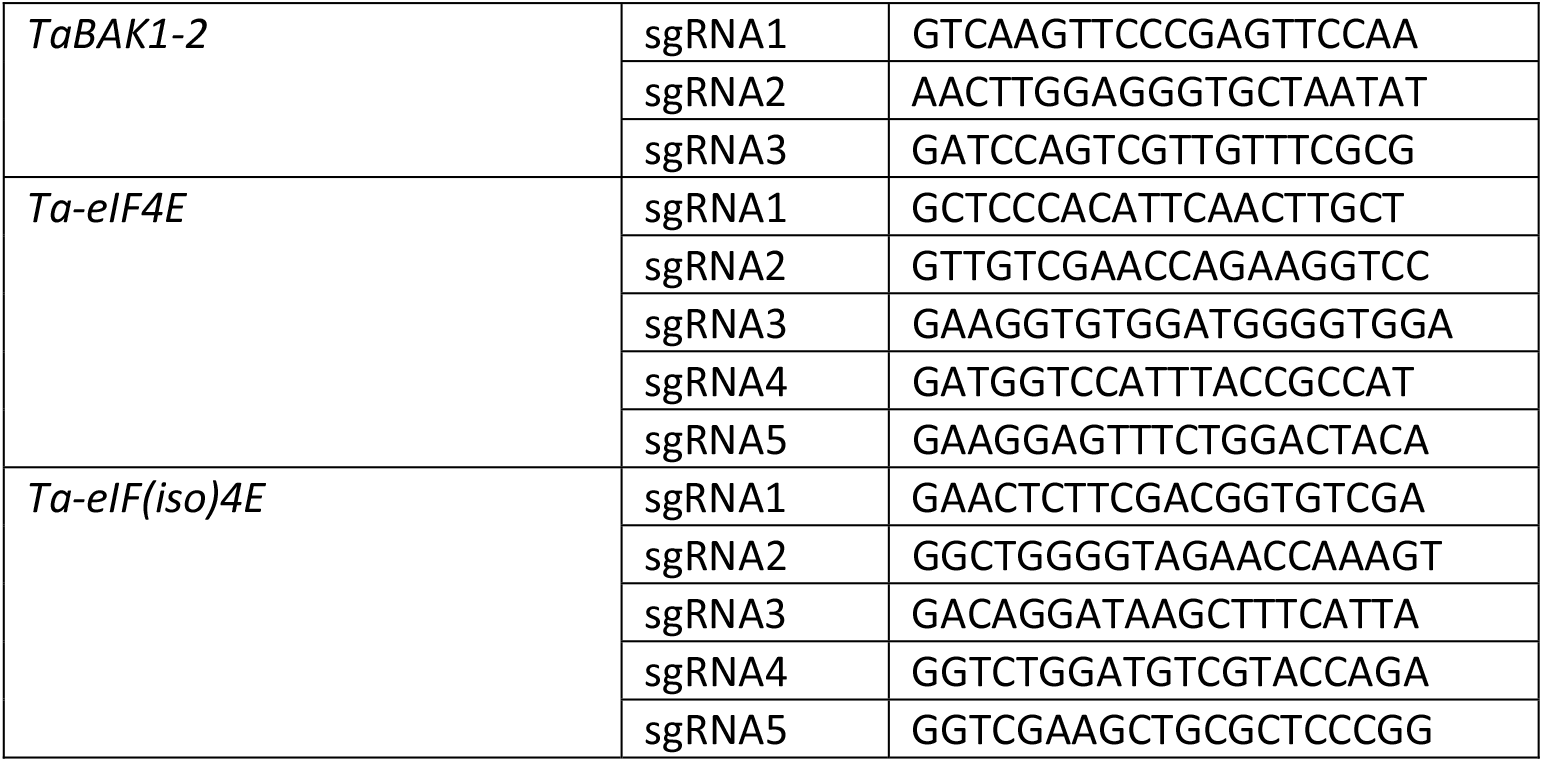
CRISPR/Cas targets.

### Plasmid construction

#### TaBAK1-2

Three guides (guides 1, 2 and 3; Figures S1-S3) targeting *TaBAK1-2* were delivered on separate constructs, not as a multiplex array. Each guide was placed under the rice U6 promoter by cloning into the pUC19_rice_sgRNA_v2 vector (kindly provided by Keith Edwards, U. of Bristol) using BtgZI as previously described for pENTR4-sgRNA4 (Zhou et al., 2014). All three sgRNA plasmids were co-delivered along with pCas9-GFP (Zhang *et al*., 2019) encoding the wheat codon-optimised Cas9, and pRRes1.111 (Alotaibi et al., 2018) encoding the *bar* selectable marker into immature wheat embryos (cv. Cadenza) as described below.

#### Ta-eIF4E/Ta-eIF(iso)4E

To express five sgRNAs per target gene, we used sgRNA-tRNA-arrays which were constructed using a modified cloning strategy based on the report by Xie et al. (2015). In all cases, we used the Gly-tRNA sequence and an improved sgRNA backbone (Dang et al., 2015).

To target *Ta-eIF4E*, six PCRs were performed using Q5 proof-reading polymerase (NEB) with vector pUC57-R504 (kindly provided by Alison Huttly, Rothamsted Research) as template and primer pairs FH187/FH188, FH189/190, FH191/192, FH193/194, FH195/196, FH197/198. Gel-extracted PCR products were assembled in a cut-ligation reaction via BsaI-HFv2 (New England Biolabs) into vector pRRES208.482 (kindly provided by Alison Huttly, Rothamsted Research) for expression under the OsU3 promoter using previously described reaction conditions (Hahn et al., 2020), resulting in vector pFH11 (Figure S4).

Similarly, to target *Ta-eIF(iso)4E*, six PCRs were performed on vector pUC57-R504 with primer pairs FH187/FH199, FH200/FH201, FH202/FH203, FH204/FH205, FH206/FH207, FH208/FH198 and the PCR amplicons were cut-ligated into pRRES208.482 via BsaI-HFv2, resulting in vector pFH12 (Figure S5).

pFH11 was combined with pFH23 (Hahn et al., 2020), encoding wheat codon-optimised Cas9 placed under the maize ubiquitin promoter (ZmUbiPr::SpCas9), and pRRes1.111 (Alotaibi et al., 2018) encoding the *bar* selectable marker. All three plasmids were co-delivered into immature wheat embryos (cvs Cezanne, Goncourt and Prevert) as described below.

pFH12 was combined with pFH23 (Hahn et al., 2020), encoding wheat codon-optimised Cas9 placed under the maize ubiquitin promoter (ZmUbiPr::SpCas9), and pRRes1.111 (Alotaibi et al., 2018) encoding the *bar* selectable marker. All three plasmids were co-delivered into immature wheat embryos (cv. Cezanne, Goncourt and Prevert) as described below.

### Growth of donor plants

The following bread wheat varieties were used for transformation: Cadenza (spring), Cezanne (winter), Goncourt (winter) and Prevert (winter).

Plants of each variety were grown in controlled environment rooms at 18°C/15°C day/night temperatures and ∼700 µM PAR for a 16 hour photoperiod. The winter varieties were initially given an 8 week vernalisation phase at 4-5°C with ∼150 µM PAR for an 8 hour photoperiod.

### Transformation

Wheat embryos of all varieties were transformed via particle bombardment essentially as previously described (Sparks and Doherty, 2020).

Donor plants were grown as above for 10-12 weeks to provide immature embryos which were isolated at 12-16 days post anthesis (dpa). The shoot/root axis was removed and the immature scutella were plated ∼ 30 per plate on the induction medium (Sparks and Doherty, 2020), and used as target tissue, giving one day pre-culture at 22°C, dark, prior to bombardment.

0.6µm gold particles (BioRad Laboratories Ltd, UK) were coated with plasmid DNAs as specified below and co-bombarded into tissues of the relevant wheat varieties using a rupture pressure of 650psi and 28.5” Hg vacuum. Following bombardment, the embryos were cultured and selected using glufosinate ammonium and putative transgenic plantlets were transferred to glasshouse conditions (all according to Sparks and Doherty, 2020).

In the case of *TaBAK1-2*, tissue culture regenerated plants were screened for the presence of transforming plasmids using the following PCR primers (see Table 2): UbiPro4 + WheatCas9R1 to test for pCas9-GFP, M13F + M13R – for sgRNA plasmids (one or more) and Bar1 + Bar2 – for pRRes1.111.

**Table 2.**
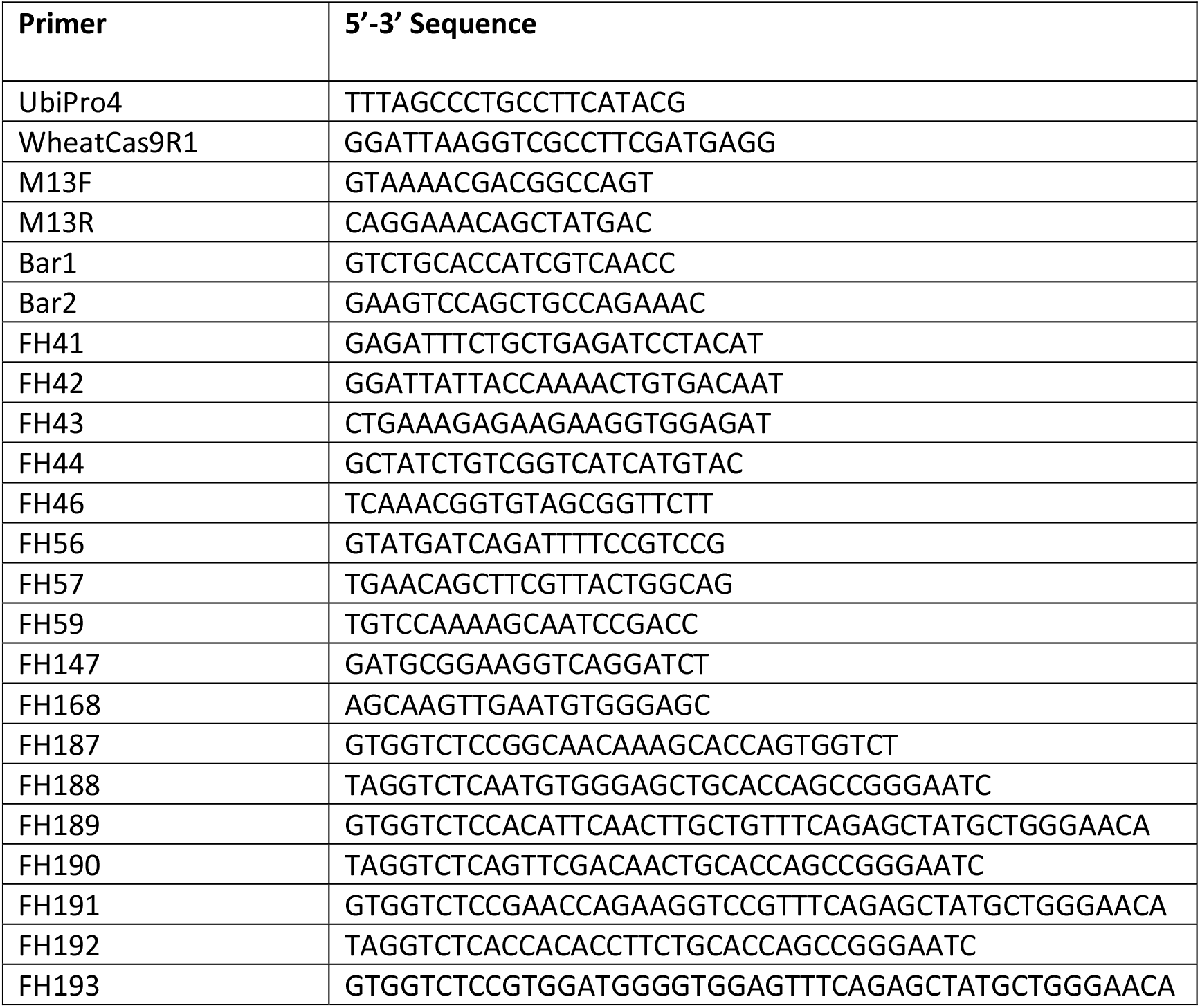
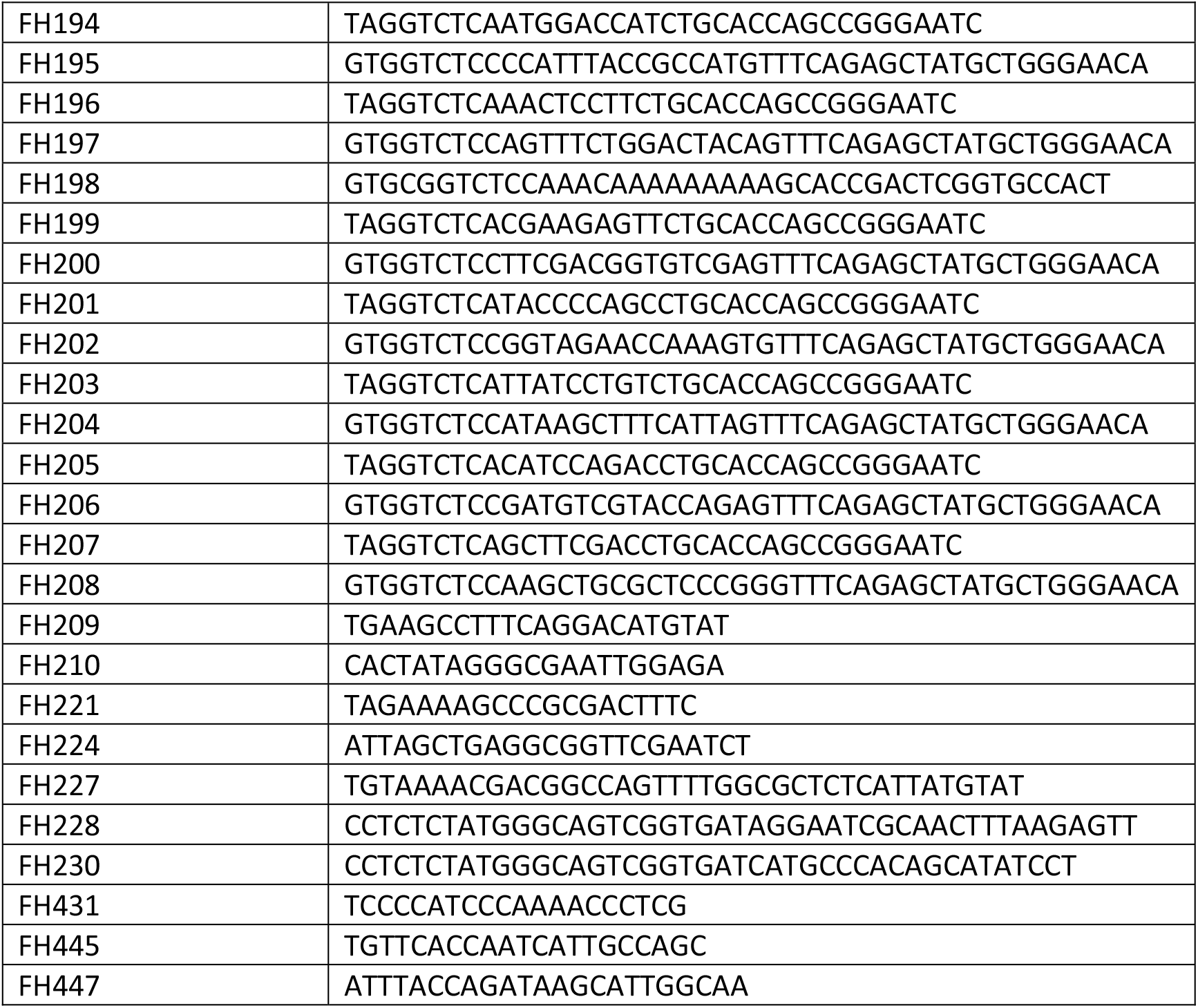
Primers used for genotyping of transgenic lines.

In the case of *Ta-eIF4E* and *Ta-eIF(iso)4E*, tissue culture regenerated plants were screened for the presence of transforming plasmids using the following PCR primers (see Table 2): UbiPro4 + FH147 to test for pFH23, FH209 + FH168 – for pFH11, FH209 + FH210 – for pFH12 and Bar1 + Bar2 – for pRRes1.111.

All plants generated from experiments were screened for CRISPR/Cas-induced indels using the PCR band shift assay (Nekrasov et al., 2017), whether PCR positive or negative for plasmids used for transformation.

### Analysis of CRISPR/Cas-induced mutations

#### TaBAK1-2

Similarly, primary (T0) transformants were analysed for mutations in the *TaBAK1-2* gene using the PCR band shift assay with primers FH41/FH44 (amplifying across all three sgRNA targets), FH41/FH42 (amplifying across sgRNA1 and 2 targets) and primers FH43/FH44 (amplifying across the sgRNA3 target). If amplicon band shifts were visible, target genes were amplified again using Q5 DNA Polymerase (New England Biolabs) with the same primer pairs as before. The PCR products were sub-cloned using the Zero Blunt™ TOPO™ PCR Cloning Kit (Thermo Fisher Scientific) and multiple single clones were Sanger-sequenced (Eurofins Genomics) to detect and analyse mutations in all subgenomes.

In the case of *TaBAK1-2*, allele profiling was performed by PCR for T1 progeny lines derived from one of the T0 transformants. This was possible because each of the *TaBAK1-2* alleles from all three subgenomes carried distinct indels that resulted in clearly distinguishable migration patterns of PCR products.

#### Ta-eIF4E and Ta-eIF(iso)4E

Primary (T0) transformants were initially screened for CRISPR/Cas-induced mutations in the *Ta-eIF4E* and *Ta-eIF(iso)4E* genes using the PCR band shift assay. For this, the target genes were amplified using DreamTaq DNA polymerase (ThermoFisher Scientific) and primers FH46 + FH221 (full gene), FH445 + FH221 (5’ part of the gene) and FH447 + FH46 (3’ part of the gene) (Table 2) in plants transformed with the vector pFH11 (in the case of targeting *Ta-eIF4E*) or primers FH431 + FH224 (full gene), FH431 + FH59 (5’ part of gene) and FH56 + FH57 (3’ part of gene) (Table 2) in plants transformed with the vector pFH12 (in the case of targeting *Ta-eIF(iso)4E*). If band shifts were detected, the respective gene fragments were amplified using Q5® DNA Polymerase and subcloned. Multiple single clones were Sanger-sequenced, as described above for *TaBAK1-2*.

## Results

### Targeted mutagenesis of the *TaBAK1-2* gene

We used the CRISPR/Cas system to target three *TaBAK1-2* homoeologues located on chromosome 2: *TraesCS2A02G343100* (*TaBAK1-2A*), *TraesCS2B01G340700* (*TaBAK1-2B*) and *TraesCS2D02G321400* (*TaBAK1-2D*). All three homoeologues were targeted at three conserved sgRNA target sites within exons 4 (sgRNA1), 5 (sgRNA2) and 11 (sgRNA3) (Figure 1A). We transformed wheat immature embryos (cv Cadenza) with DNA constructs expressing CRISPR/Cas reagents, as described in Materials and Methods, and regenerated 30 T0 primary transgenic lines. We then genotyped the T0 lines for the presence of CRISPR/Cas-induced indels using the PCR band shift assay (Nekrasov et al., 2017). Out of the 30 T0 lines, two showed a clear PCR band shift indicating presence of deletions of around 600 bp (Figure 1B). To identify *TaBAK1-2* alleles present in both T0 plants, we subcloned the PCR amplicons into a high copy number vector and sequenced individual clones by Sanger. The T0 plant 1 turned out to be a triple-biallelic carrying indels in all six copies of *TaBAK1*, while the plant 2 carried heterozygous mutations in *TaBAK1-2A* and *TaBAK1-2D* and no mutations in *TaBAK1-2B* (Figure 1C and D). The three insertions identified in the T0 plant 1 (43 bp, 81 bp and 136 bp; Figure 1C) proved to be fragments of the pCas9-GFP plasmid. Such insertion events at CRISPR/Cas target sites were previously reported in potato (Andersson et al., 2017, 2018).

**Figure 1.**
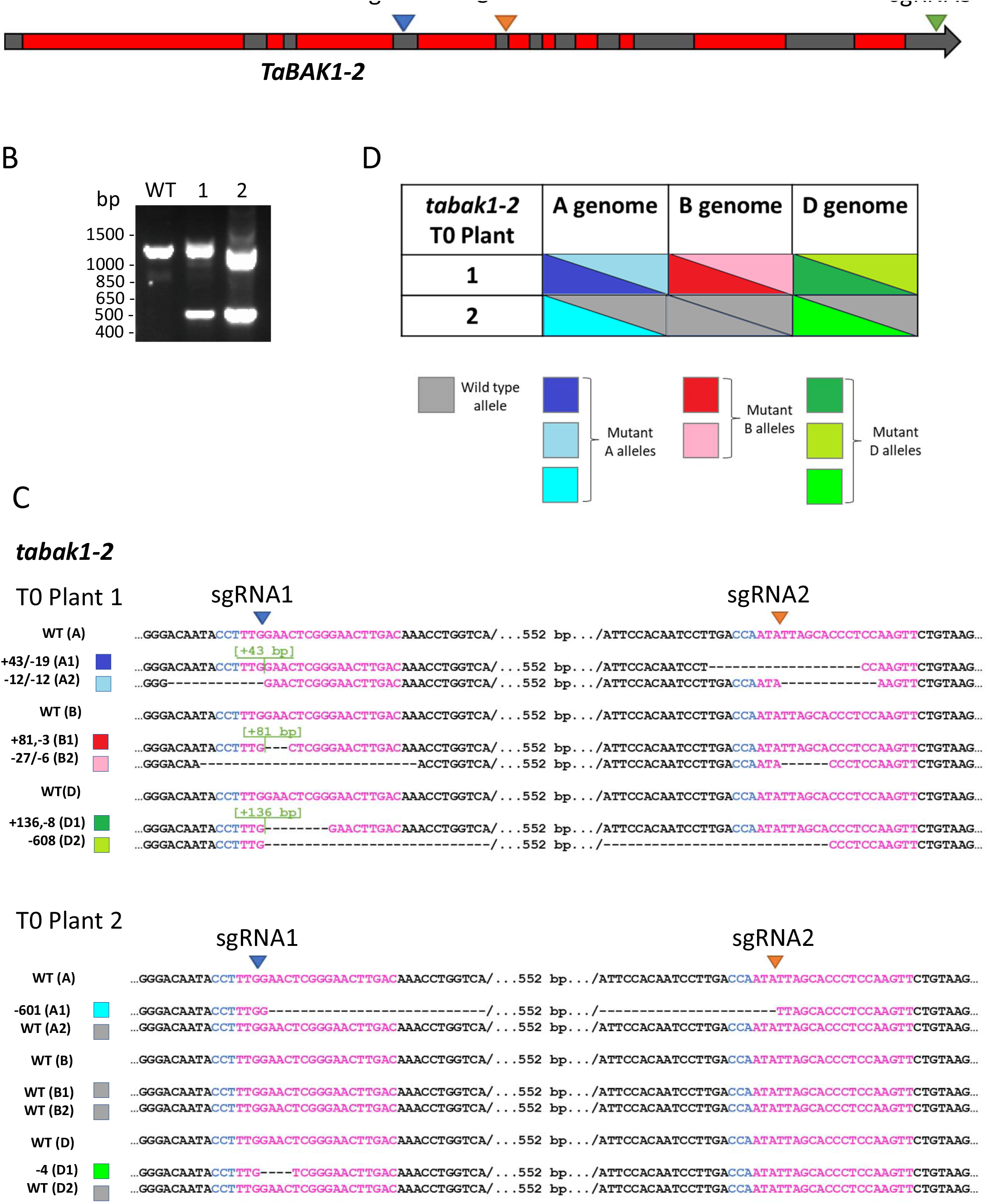
Targeted mutagenesis of *TaBak1-2*. (A) Cartoon showing locations of sgRNA target sites. (B) PCR genotyping of CRISPR/Cas-mutagenised T0 lines. (C) Mutant *tabak1-2* alleles identified by Sanger sequencing in T0 plants. (D) Allele composition of *tabak1-2* T0 plants.

It should be noted that PCR amplicons shown in Figure 1B were produced using primers amplifying across the sgRNA1 and 2 target sites. We also separately amplified across the sgRNA3 target in both T0 plants and did not detect any mutations at this site.

We selected the *tabak1-2* T0 line 1 for further analysis as this line showed mutations in all six copies of the *TaBAK1-2* gene.

To check if the mutations present in the *tabak1-2* T0 line 1 could be transmitted through the germline and inherited by the next generation, we PCR-genotyped 52 T1 progeny plants derived from this line. The genotyping data clearly indicated inheritance of all six mutant alleles (Figures 2, S6 and S7). Out of 52 T1 lines, five were triple-homozygous (plants 4, 19, 23, 43 and 52; Figure S7).

**Figure 2.**
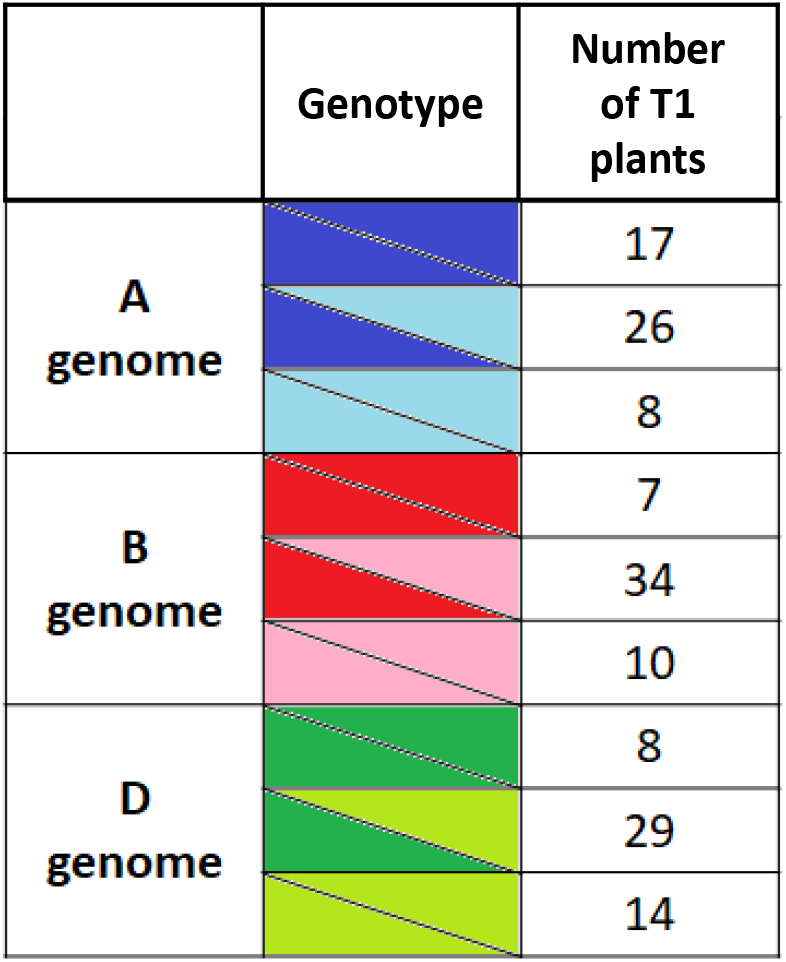
CRISPR/Cas-induced *tabak1-2* mutant alleles are inherited by the next generation. The table shows allele distribution among T1 progeny of the T0 plant 1.

### Targeted mutagenesis of the *Ta-eIF4E* and *Ta-eIF(iso)4E* genes

We targeted both *Ta-eIF4E* (homoeologues *TraesCS3A02G521500, TraesCS3B02G591300* and *TraesCS3D02G527800*) and *Ta-eIF(iso)4E* (homoeologues *TraesCS1A02G149200, TraesCS1B02G167100* and *TraesCS1D02G146500*) genes with five sgRNAs each (Figures 3A and 4A, respectively) in three winter cultivars of common wheat, Cezanne, Goncourt and Prevert, which are susceptible to WSSMV (Dragan Perovic, personal communication).

**Figure 3.**
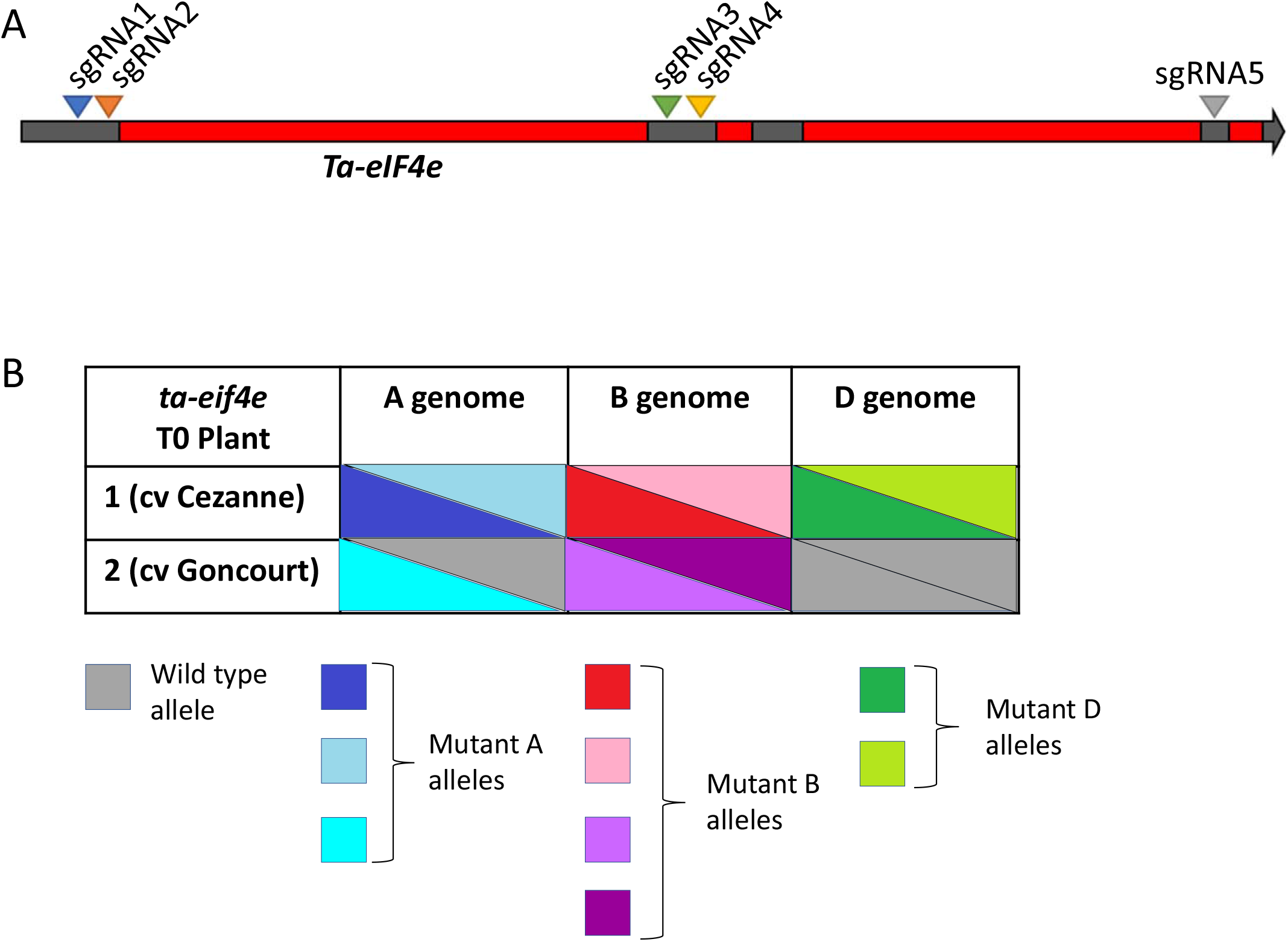
Targeted mutagenesis of *Ta-eIF4e*. (A) Cartoon showing locations of sgRNA target sites. (B) Allele composition of *ta-eif4e* T0 plants.

In total, we screened 49 T0 plants for *Ta-eIF4E* (40, 8 and 1 from cvs Cezanne, Goncourt and Prevert transformations, respectively) and 40 T0 plants for *Ta-eIF(iso)4E* (30, 5 and 5 from cvs Cezanne, Goncourt and Prevert transformations, respectively). Genotyping identified two T0 plants carrying CRISPR/Cas-induced indels in *Ta-eIF4E* (cvs Cezanne and Goncourt; Figures 3B and S8) and two T0 plants with indels in *Ta-eIF(iso)4E* (cvs Cezanne and Prevert; Figures 4B, S9 and S10). Two out of the four T0 plants were triple-biallelic: *ta-eif4e* T0 plant 1 (cv Cezanne; Figures 3B and S8) and *ta-eif(iso)4e* T0 plant 2 (cv Prevert; Figures 4B and S10).

**Figure 4.**
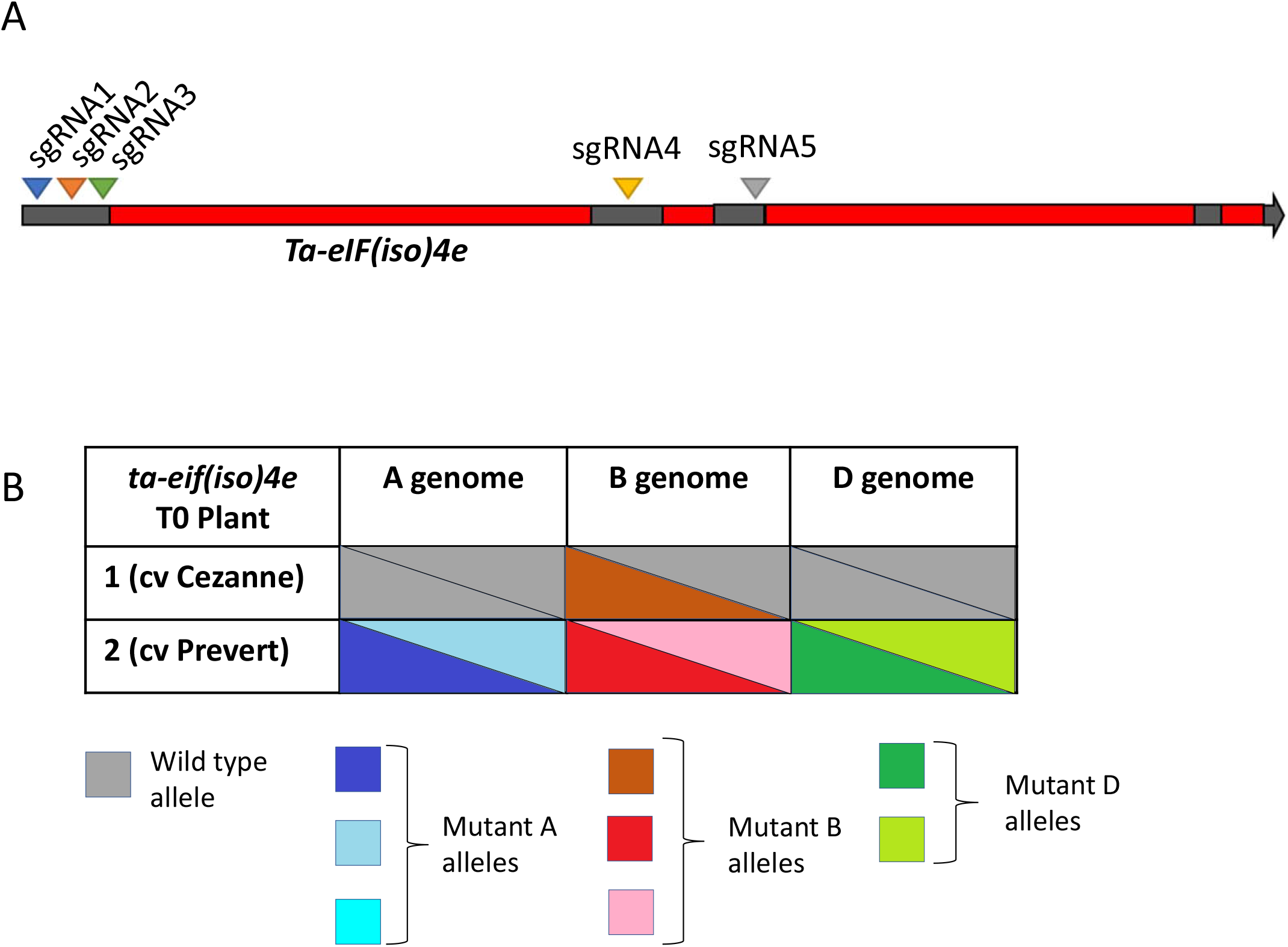
Targeted mutagenesis of *Ta-eIF(iso)4e*. (A) Cartoon showing locations of sgRNA target sites. (B) Allele composition of *ta-eif(iso)4e* T0 plants.

As in the case of *TaBAK1-2*, we detected CRISPR/Cas-induced insertions in *ta-eif4e* and *ta-eif(iso)4e* T0 lines ranging from 242 bp to 592 bp (Figures S8-S10). The inserted DNA was derived from plasmids used for transformation, wheat genomic DNA, bacterial DNA or combinations of those.

## Discussion

During this study we generated *tabak1-2* lines carrying different combinations of mutant *tabak1-2a, tabak1-2b and tabak1-2d* alleles (Figure S7). BAK1 acts as a coreceptor for a number of pattern recognition receptors (PRRs) involved in pattern-triggered immunity (PTI) responses in plants (Yasuda et al., 2017). The generated *tabak1-2* alleles are expected to be loss-of-function as they carry rather large deletions/insertions within the coding regions or short deletions (e.g. -4 bp) that put the coding sequence out of frame. We therefore expect the *tabak1-2* lines to become a useful genetic resource for the research community working on molecular mechanisms of plant-microbe interactions in wheat.

To generate the *ta-eif4e* and *ta-eif(iso)4e* lines, we needed to optimise the transformation procedure for the three winter wheat cultivars Cezanne, Goncourt and Prevert (see Materials and Methods). Since there are only few published examples of genome editing in winter wheat (Li et al., 2020; Liu et al., 2021), our work will be of interest to researchers working on winter wheat transformation and genome editing.

To be able to evaluate *ta-eif4e* and *ta-eif(iso)4e* mutants for enhanced resistance to WSSMV, phenotypic characterisation of these lines would need to be carried out in a follow-up study. Since WSSMV is transmitted by *Polymyxa graminis* (Jiang et al., 2020), a soil-borne filamentous microorganism infecting wheat roots, it is very difficult to perform WSSMV pathotests under the glasshouse conditions. On the other hand, GM field trials in the south of Europe, where WSSMV can be found, are problematic right now due to the policy restrictions and significant public opposition. We are hopeful the situation will change at some point in the future. It should be noted, that since some of the *ta-eif4e* and *ta-eif(iso)4e* mutant alleles contain inserted fragments of transgenic or wheat genomic DNA in them, plants carrying them cannot be treated as transgene-free genome edited but rather GM lines.

## Acknowledgements

We thank Angela Doherty, Melloney St-Leger, Andrey Korolev and Lucy Hyde for excellent technical assistance.

We thank Keith Edwards (University of Bristol, UK) for pCas9-GFP and pUC19_rice_sgRNA_v2, and Alison Huttly (Rothamsted Research, UK) for pUC57-R504 and pRRES208.482 DNA constructs. We thank Dragan Perovic (Julius Kühn-Institute, Germany) for advice on choosing wheat cultivars for CRISPR/Cas mutagenesis of *Ta-eIF4E* and *Ta-eIF(iso)4E* genes and providing the seed of Cezanne, Goncourt and Prevert wheat varieties.

This research was supported by the Biotechnology and Biological Sciences Research Council (BBSRC) through the Designing Future Wheat (DFW) Institute Strategic Programme.

