## Supplementary Figures for "Efficient CRISPR/Cas-mediated Targeted Mutagenesis in Spring and Winter Wheat Varieties"

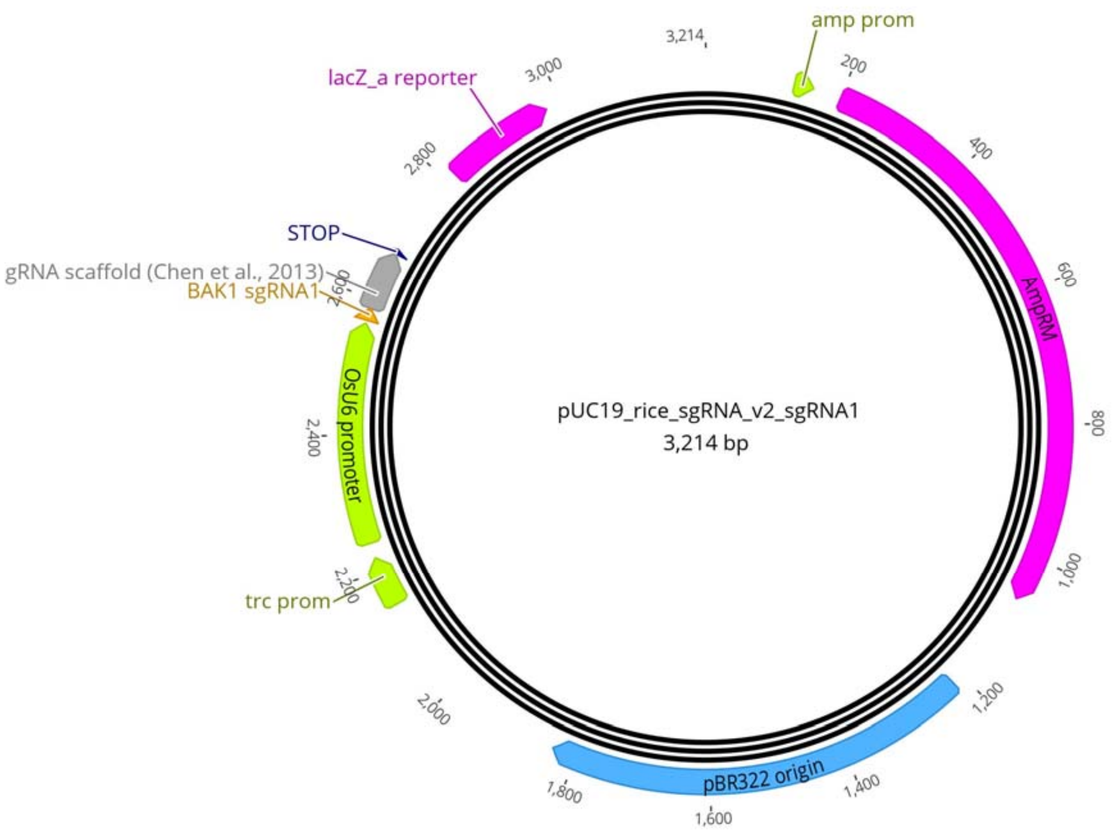


AAGAACGAACTAAGCCGGACAAAAAAAGGAGCACATATACAAACCGGTTTTATTCATGAATGGTCACGATGGATGATGG
GGCTCAGACTTGAGCTACGAGGCCGCAGGCGAGAGAAGCCTAGTGTGCTCTCTGCTTGTTTGGGCCGTAACGGAGGATA
CGGCCGACGAGCGTGTACTACCGCGCGGGATGCCGCTGGGCGCTGCGGGGGCCGTTGGATGGGGATCGGTGGGTCGCGG
GAGCGTTGAGGGGAGACAGGTTTAGTACCACCTCGCCTACCGAACAATGAAGAACCCACCTTATAACCCCGCGCGCTGC
CGCTTGTGTTGTCAAGTTCCCGAGTTCCAAGTTTAAGAGCTATGCTGGAAACAGCATAGCAAGTTTAAATAAGGCTAGT
CCGTTATCAACTTGAAAAAGTGGCACCGAGTCGGTGCTTTTTTTT

OsU6 promoter
gRNA scaffold (Chen et al., 2013)
PolyT STOP
Target sequence

**Figure S1**
pUC19_rice_sgRNA_v2_sgRNA1 construct map and insert sequence.

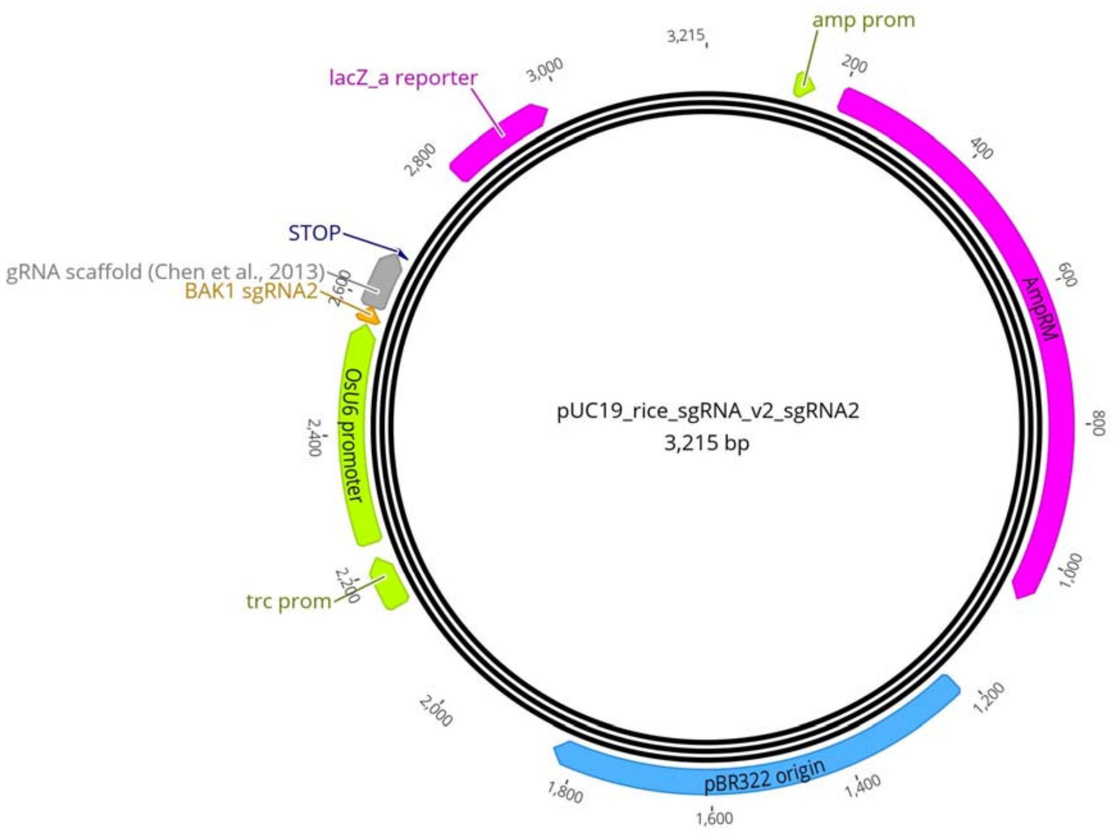


AAGAACGAACTAAGCCGGACAAAAAAAGGAGCACATATACAAACCGGTTTTATTCATGAATGGTCACGATGGATGATGG
GGCTCAGACTTGAGCTACGAGGCCGCAGGCGAGAGAAGCCTAGTGTGCTCTCTGCTTGTTTGGGCCGTAACGGAGGATA
CGGCCGACGAGCGTGTACTACCGCGCGGGATGCCGCTGGGCGCTGCGGGGGCCGTTGGATGGGGATCGGTGGGTCGCGG
GAGCGTTGAGGGGAGACAGGTTTAGTACCACCTCGCCTACCGAACAATGAAGAACCCACCTTATAACCCCGCGCGCTGC
CGCTTGTGTTGAACTTGGAGGGTGCTAATATGTTTAAGAGCTATGCTGGAAACAGCATAGCAAGTTTAAATAAGGCTAG
TCCGTTATCAACTTGAAAAAGTGGCACCGAGTCGGTGCTTTTTTTT

OsU6 promoter
gRNA scaffold (Chen et al., 2013)
PolyT STOP
Target sequence

**Figure S2**
pUC19_rice_sgRNA_v2_sgRNA2 construct map and insert sequence.

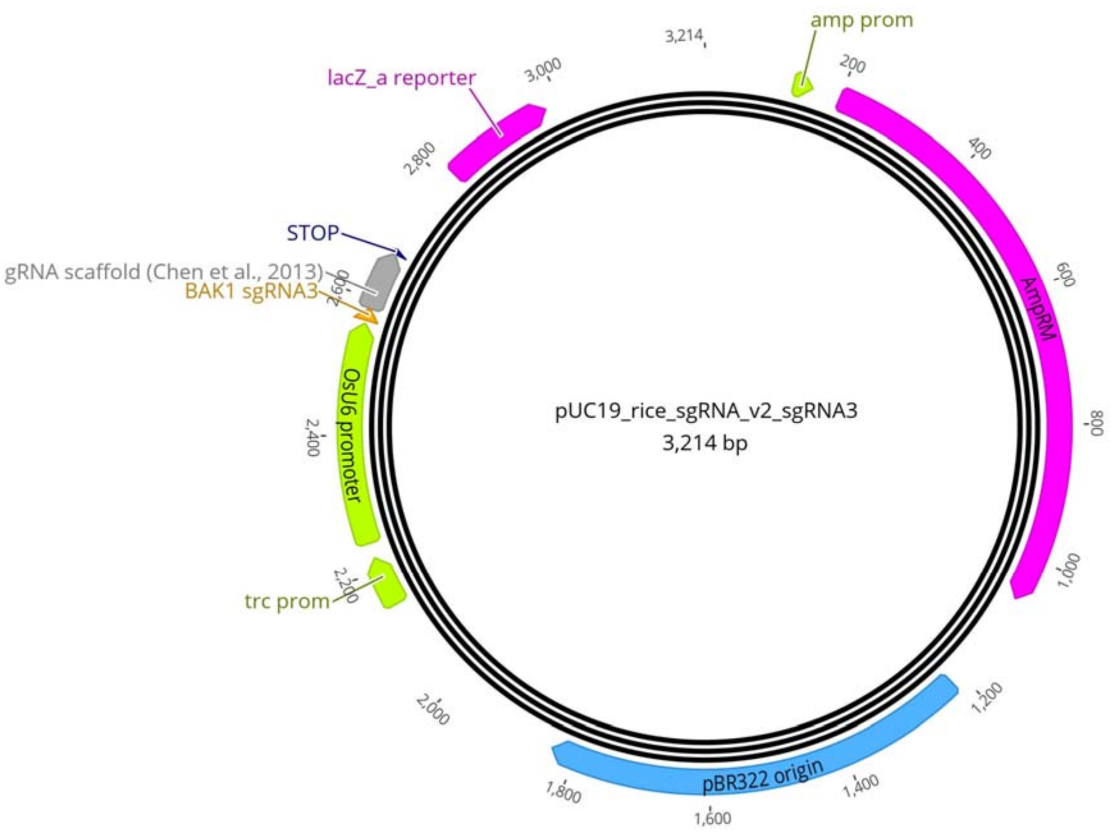


AAGAACGAACTAAGCCGGACAAAAAAAGGAGCACATATACAAACCGGTTTTATTCATGAATGGTCACGATGGATGATGG
GGCTCAGACTTGAGCTACGAGGCCGCAGGCGAGAGAAGCCTAGTGTGCTCTCTGCTTGTTTGGGCCGTAACGGAGGATA
CGGCCGACGAGCGTGTACTACCGCGCGGGATGCCGCTGGGCGCTGCGGGGGCCGTTGGATGGGGATCGGTGGGTCGCGG
GAGCGTTGAGGGGAGACAGGTTTAGTACCACCTCGCCTACCGAACAATGAAGAACCCACCTTATAACCCCGCGCGCTGC
CGCTTGTGTTGATCCAGTCGTTGTTTCGCGGTTTAAGAGCTATGCTGGAAACAGCATAGCAAGTTTAAATAAGGCTAGT
CCGTTATCAACTTGAAAAAGTGGCACCGAGTCGGTGCTTTTTTTT

OsU6 promoter
gRNA scaffold (Chen et al., 2013)
PolyT STOP
Target sequence

**Figure S3**
pUC19_rice_sgRNA_v2_sgRNA3 construct map and insert sequence.

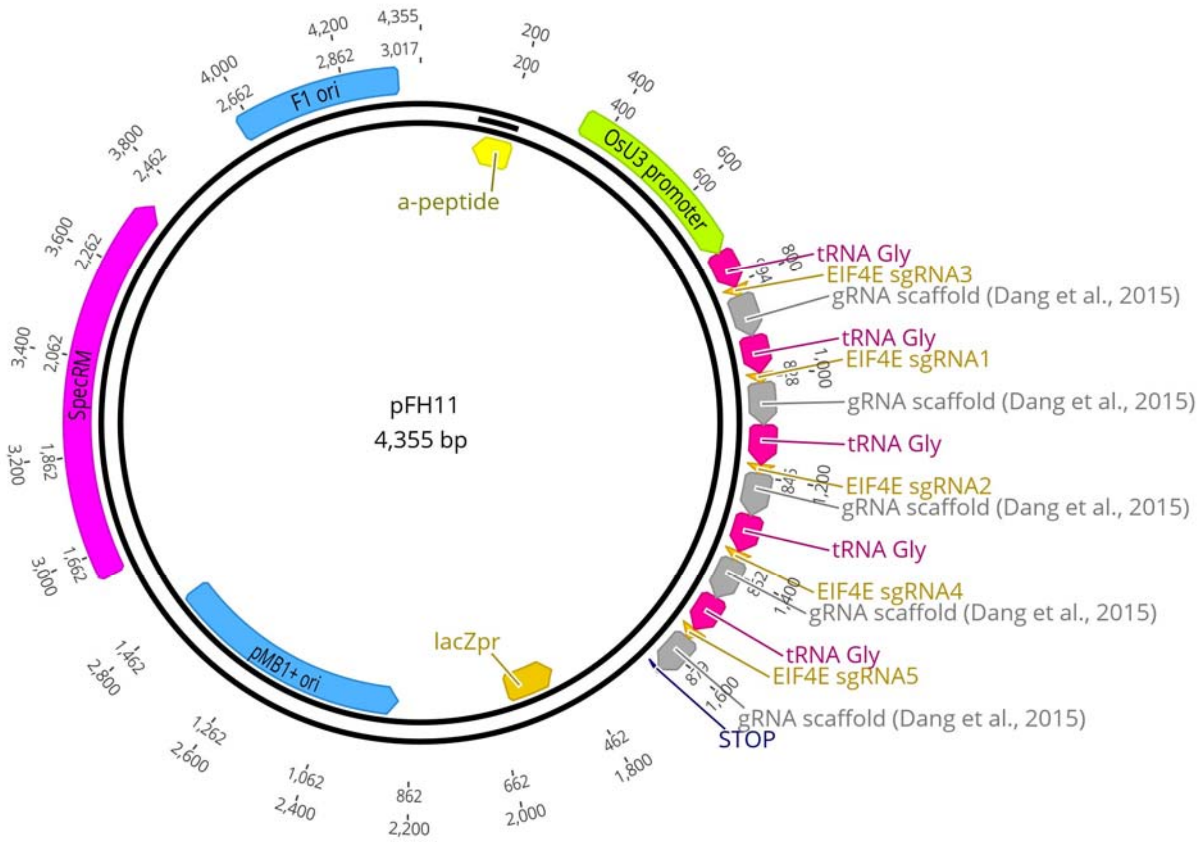


AAGCTTAAGGAATCTTTAAACATACGAACAGATCACTTAAAGTTCTTCTGAAGCAACTTAAAGTTATCAGGCATGCATG
GATCTTGGAGGAATCAGATGTGCAGTCAGGGACCATAGCACAAGACAGGCGTCTTCTACTGGTGCTACCAGCAAATGCT
GGAAGCCGGGAACACTGGGTACGTTGGAAACCACGTGATGTGAAGAAGTAAGATAAACTGTAGGAGAAAAGCATTTCGT
AGTGGGCCATGAAGCCTTTCAGGACATGTATTGCAGTATGGGCCGGCCCATTACGCAATTGGACGACAACAAAGACTAG
TATTAGTACCACCTCGGCTATCCACATAGATCAAAGCTGATTTAAAAGAGTTGTGCAGATGATCCGTGGCAACAAAGCA
CCAGTGGTCTAGTGGTAGAATAGTACCCTGCCACGGTACAGACCCGGGTTCGATTCCCGGCTGGTGCAGCTCCCACATT
CAACTTGCTGTTTCAGAGCTATGCTGGGAACAGCATAGCAAGTTGAAATAAGGCTAGTCCGTTATCAACTTGAAAAAGT
GGCACCGAGTCGGTGCAACAAAGCACCAGTGGTCTAGTGGTAGAATAGTACCCTGCCACGGTACAGACCCGGGTTCGAT
TCCCGGCTGGTGCAGTTGTCGAACCAGAAGGTCCGTTTCAGAGCTATGCTGGGAACAGCATAGCAAGTTGAAATAAGGC
TAGTCCGTTATCAACTTGAAAAAGTGGCACCGAGTCGGTGCAACAAAGCACCAGTGGTCTAGTGGTAGAATAGTACCCT
GCCACGGTACAGACCCGGGTTCGATTCCCGGCTGGTGCAGAAGGTGTGGATGGGGTGGAGTTTCAGAGCTATGCTGGGA
ACAGCATAGCAAGTTGAAATAAGGCTAGTCCGTTATCAACTTGAAAAAGTGGCACCGAGTCGGTGCAACAAAGCACCAG
TGGTCTAGTGGTAGAATAGTACCCTGCCACGGTACAGACCCGGGTTCGATTCCCGGCTGGTGCAGATGGTCCATTTACC
GCCATGTTTCAGAGCTATGCTGGGAACAGCATAGCAAGTTGAAATAAGGCTAGTCCGTTATCAACTTGAAAAAGTGGCA
CCGAGTCGGTGCAACAAAGCACCAGTGGTCTAGTGGTAGAATAGTACCCTGCCACGGTACAGACCCGGGTTCGATTCCC
GGCTGGTGCAGAAGGAGTTTCTGGACTACAGTTTCAGAGCTATGCTGGGAACAGCATAGCAAGTTGAAATAAGGCTAGT
CCGTTATCAACTTGAAAAAGTGGCACCGAGTCGGTGCTTTTTTTTT

OsU3 promoter
tRNA Gly
gRNA scaffold (Dang et al., 2015)
PolyT STOP
Target sequences

**Figure S4**
pFH11 construct map and insert sequence.

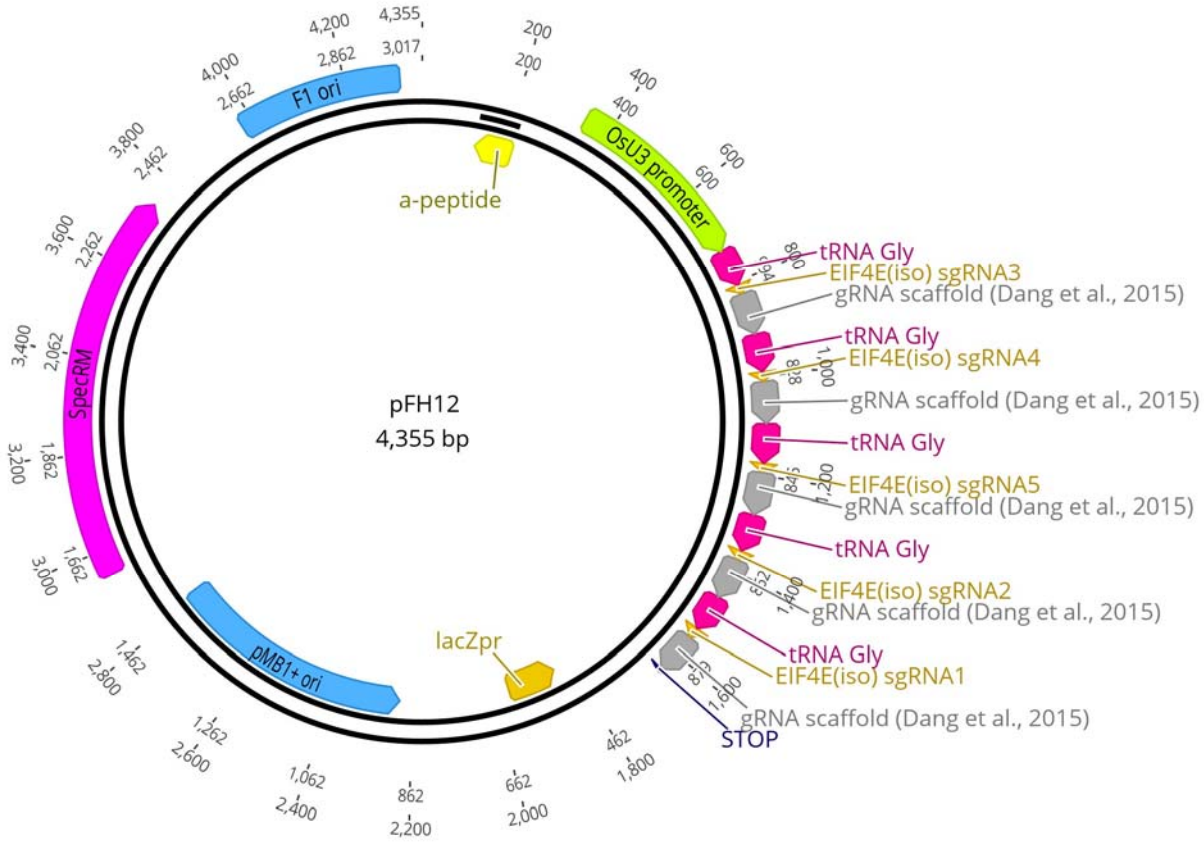


AAGCTTAAGGAATCTTTAAACATACGAACAGATCACTTAAAGTTCTTCTGAAGCAACTTAAAGTTATCAGGCATGCATG
GATCTTGGAGGAATCAGATGTGCAGTCAGGGACCATAGCACAAGACAGGCGTCTTCTACTGGTGCTACCAGCAAATGCT
GGAAGCCGGGAACACTGGGTACGTTGGAAACCACGTGATGTGAAGAAGTAAGATAAACTGTAGGAGAAAAGCATTTCGT
AGTGGGCCATGAAGCCTTTCAGGACATGTATTGCAGTATGGGCCGGCCCATTACGCAATTGGACGACAACAAAGACTAG
TATTAGTACCACCTCGGCTATCCACATAGATCAAAGCTGATTTAAAAGAGTTGTGCAGATGATCCGTGGCAACAAAGCA
CCAGTGGTCTAGTGGTAGAATAGTACCCTGCCACGGTACAGACCCGGGTTCGATTCCCGGCTGGTGCAGAACTCTTCGA
CGGTGTCGAGTTTCAGAGCTATGCTGGGAACAGCATAGCAAGTTGAAATAAGGCTAGTCCGTTATCAACTTGAAAAAGT
GGCACCGAGTCGGTGCAACAAAGCACCAGTGGTCTAGTGGTAGAATAGTACCCTGCCACGGTACAGACCCGGGTTCGAT
TCCCGGCTGGTGCAGGCTGGGGTAGAACCAAAGTGTTTCAGAGCTATGCTGGGAACAGCATAGCAAGTTGAAATAAGGC
TAGTCCGTTATCAACTTGAAAAAGTGGCACCGAGTCGGTGCAACAAAGCACCAGTGGTCTAGTGGTAGAATAGTACCCT
GCCACGGTACAGACCCGGGTTCGATTCCCGGCTGGTGCAGACAGGATAAGCTTTCATTAGTTTCAGAGCTATGCTGGGA
ACAGCATAGCAAGTTGAAATAAGGCTAGTCCGTTATCAACTTGAAAAAGTGGCACCGAGTCGGTGCAACAAAGCACCAG
TGGTCTAGTGGTAGAATAGTACCCTGCCACGGTACAGACCCGGGTTCGATTCCCGGCTGGTGCAGGTCTGGATGTCGTA
CCAGAGTTTCAGAGCTATGCTGGGAACAGCATAGCAAGTTGAAATAAGGCTAGTCCGTTATCAACTTGAAAAAGTGGCA
CCGAGTCGGTGCAACAAAGCACCAGTGGTCTAGTGGTAGAATAGTACCCTGCCACGGTACAGACCCGGGTTCGATTCCC
GGCTGGTGCAGGTCGAAGCTGCGCTCCCGGGTTTCAGAGCTATGCTGGGAACAGCATAGCAAGTTGAAATAAGGCTAGT
CCGTTATCAACTTGAAAAAGTGGCACCGAGTCGGTGCTTTTTTTTT

OsU3 promoter
tRNA Gly
gRNA scaffold (Dang et al., 2015)
PolyT STOP
Target sequences

**Figure S5**
pFH12 construct map and insert sequence.

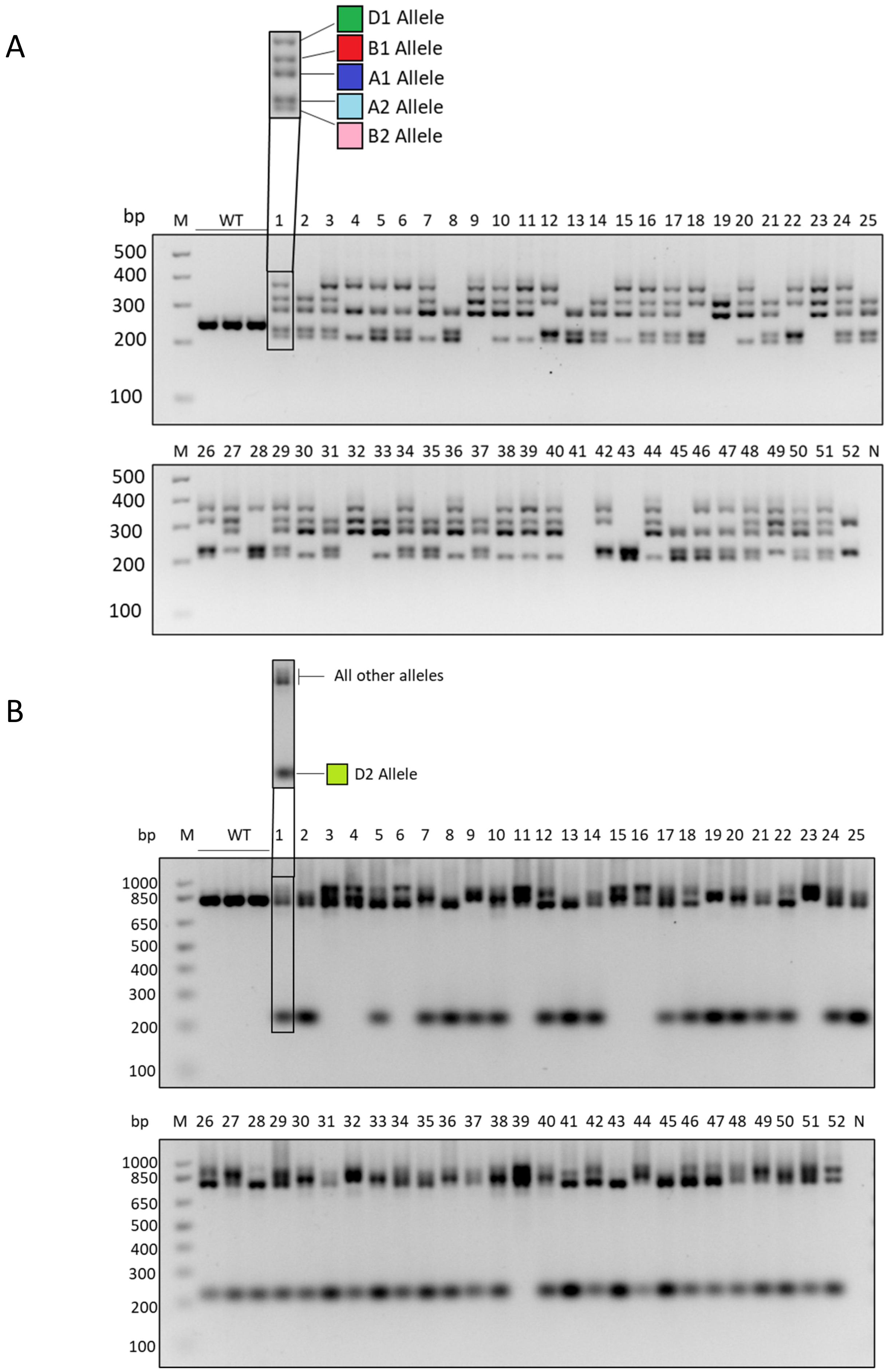


**Figure S6**

PCR-genotyping of T1 progeny of *tabak1-2* T0 plant 1 with FH227 + FH228 primers (A) and FH227 + FH230 primers (B).

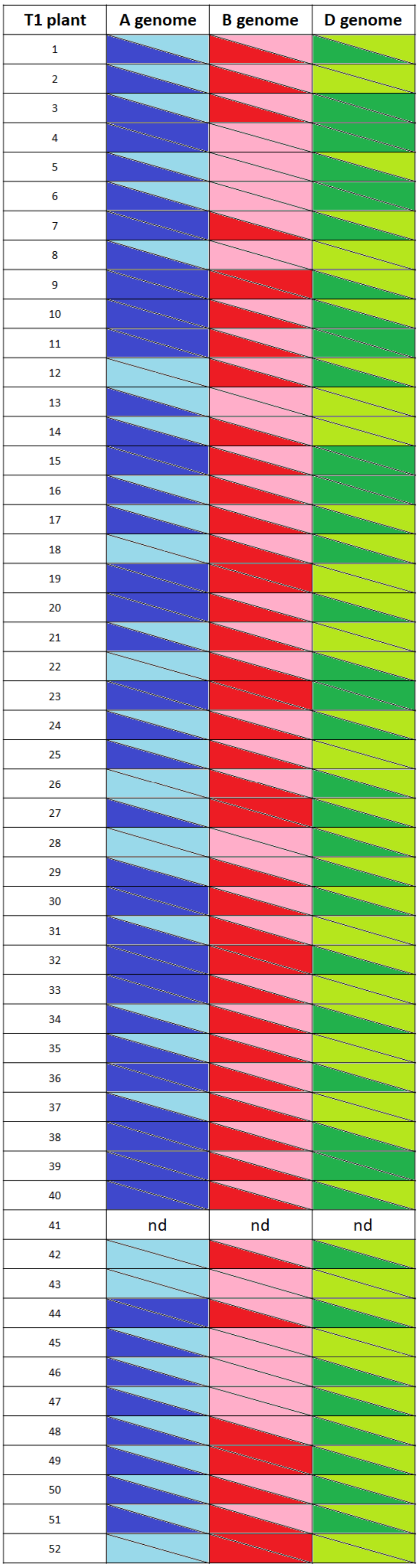


**Figure S7**

*TaBAK1-2* allele distribution among T1 progeny of *tabak1-2* T0 plant 1.

#### *ta-eif4e*

T0 Plant 1 (cv Cezanne)

```
                      sgRNA1                                                   sgRNA2
WT (A)                ...AACGCCTGGACCTTCTGGTTCGACAACCCGCAGGGC/..19 bp../GGGGGAGCACCATCCACCCCATCCACACCTTCTCCAC...

+1(A1)                ...AACGCCTGGACCTTCTGGTTCGACAACCCGCAGGGC/..19 bp../GGGGGAGCACCATCCACCCCATCCACACCTTCTCCAC...
+430, -7 (A2)         ...AACGCCTGGACCTTCTGGTTCGACAACCCGCAGGGC/..19 bp../GGGGGAGCACCATCCACCCCATCCACACCTTCTCCAC...

WT (B)                ...AACGCCTGGACCTTCTGGTTCGACAACCCGCAGGGC/..19 bp../GGGGCAGCACCATCCACCCCATCCACACCTTCTCCAC...
+1 (B1)               ...AACGCCTGGACCTTCTGGTTCGACAACCCGCAGGGC/..19 bp../GGGGCAGCACCATCCACCCCATCCACACCTTCTCCAC...
                                                                                  [+517 bp]
+517, -30 (B2)        ...AACGCCTGGACCTTCTGGTTCGACAACCCGCAGGGC/..19 bp../GGGGCAGCACCATCCACCCCATCCACACCTTCTCCAC...

WT (D)                ...AACGCCTGGACCTTCTGGTTCGACAACCCGCAGGGC/..19 bp../GGGGGAGCACCATCCACCCCATCCACACCTTCTCCAC...

+1 (D1)               ...AACGCCTGGACCTTCTGGTTCGACAACCCGCAGGGC/..19 bp../GGGGGAGCACCATCCACCCCATCCACACCTTCTCCAC...
-58 (D2)              ...AACGCCTGGACCTTCTGGTTCGACAACCCGCAGGGC/..19 bp../GGGGGAGCACCATCCACCCCATCCACACCTTCTCCAC...
```

```
                                                          sgRNA5
WT (A)                ...AGATAAGCATTGGCAAGCAGTGGAAGGAGTTTCTGGACT-ACAAGGACTCCATTGGGTTCATTGTTCATGTGAGTCGCTTGC...

+1(A1)                ...AGATAAGCATTGGCAAGCAGTGGAAGGAGTTTCTGGACTTACAAGGACTCCATTGGGTTCATTGTTCATGTGAGTCGCTTGC...
                                                          [+430 bp]
+430, -7 (A2)         ...AGATAAGCATTGGCAAGCAGTGGAAGGAGTTTCTGGAC--------ACTCCATTGGGTTCATTGTTCATGTGAGTCGCTTGC...

WT (B)                ...AGATAAGCATTGGCAAGCAGTGGAAGGAGTTTCTGGACT-ACAGGGACTCCATTGGGTTCATTGTTCATGTGAGTCGCTTGC...

+1 (B1)               ...AGATAAGCATTGGCAAGCAGTGGAAGGAGTTTCTGGACTTACAGGGACTCCATTGGGTTCATTGTTCATGTGAGTCGCTTGC...
+517, -30 (B2)        ...AGATAAGCATTGGCAAGCAGTGGAAGGAGTTTCTGG-------------------------------ATGTGAGTCGCTTGC...

WT (D)                ...AGATAAGCATTGGCAAGCAGTGGAAGGAGTTTCTGGACT-ACAAGGACTCCATTGGGTTCATTGTTCATGTGAGTCGCTTGC...

+1 (D1)               ...AGATAAGCATTGGCAAGCAGTGGAAGGAGTTTCTGGACTTACAAGGACTCCATTGGGTTCATTGTTCATGTGAGTCGCTTGC...
-58 (D2)              ...AGATAAGCAT-----------------------------------------------------------GTGAGTCGCTTGC...
```

T0 Plant 2 (cv Goncourt)

```
                                                                  sgRNA5
WT (A)                ...TTATTTACCAGATAAGCATTGGCAAGCAGTGGAAGGAGTTTCTGGACTACAAGGACTCCATTGGGTTCATTGTTCATGTGAG...
                                                                  [+592 bp]
+592, -8 (A2)         ...TTATTTACCAGATAAGCATTGGCAAGCAGTGGAAGGAGTTTCTGGA--------ACTCCATTGGGTTCATTGTTCATGTGAG...
WT (A1)               ...TTATTTACCAGATAAGCATTGGCAAGCAGTGGAAGGAGTTTCTGGACTACAAGGACTCCATTGGGTTCATTGTTCATGTGAG...

WT (B)                ...TTATTTACCAGATAAGCATTGGCAAGCAGTGGAAGGAGTTTCTGGACTACAGGGACTCCATTGGGTTCATTGTTCATGTGAG...

-66 (B1)              ...TTATTTA------------------------------------------------------------------TCATGTGAG...
-21 (B2)              ...TTATTTACCAGATAAGCATTGGCAAGCAGTGGAAGGAGTTT---------------------GGGTTCATTGTTCATGTGAG...

WT (D)                ...TTATTTACCAGATAAGCATTGGCAAGCAGTGGAAGGAGTTTCTGGACTACAAGGACTCCATTGGGTTCATTGTTCATGTGAG...

WT (D1)               ...TTATTTACCAGATAAGCATTGGCAAGCAGTGGAAGGAGTTTCTGGACTACAAGGACTCCATTGGGTTCATTGTTCATGTGAG...
WT (D2)               ...TTATTTACCAGATAAGCATTGGCAAGCAGTGGAAGGAGTTTCTGGACTACAAGGACTCCATTGGGTTCATTGTTCATGTGAG...
```

**Figure S8**
Alignments showing CRISPR/Cas-induced indels in *Ta-eIF4e* homoeologues in two *ta-eif4e* T0 plants (cvs Cezanne and Goncourt).

### *ta-eif(iso)4e*

T0 Plant 1 (cv Cezanne)

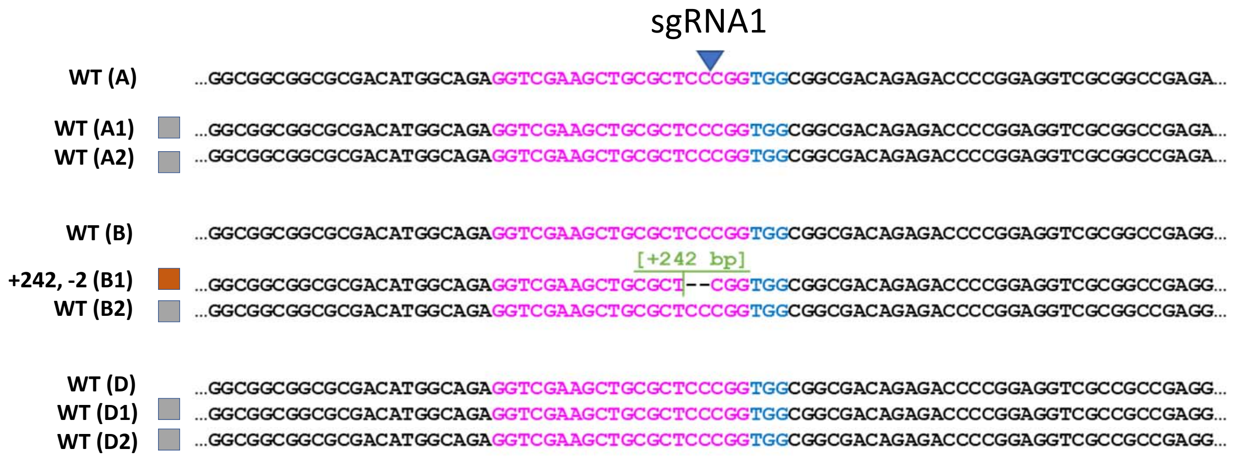


**Figure S9**

Alignment showing CRISPR/Cas-induced indels in *Ta-eIF(iso)4e* homoeologues in the *ta-eif(iso)4e* T0 plant 1 (cv Cezanne).

### *ta-eIf(iso)4e*

T0 Plant 2 (cv Prevert)

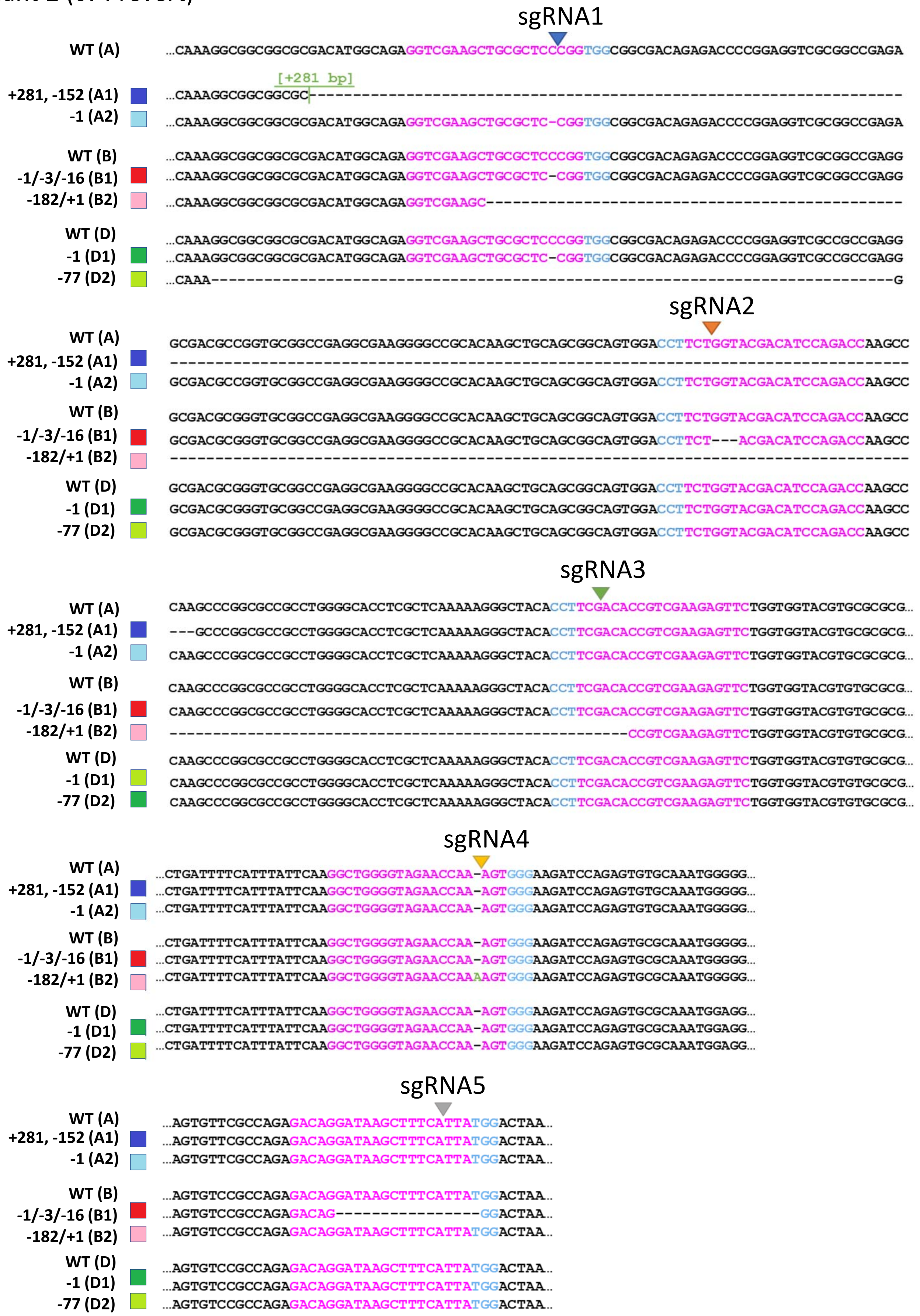


**Figure S10**

Alignment showing CRISPR/Cas-induced indels in *Ta-eIF(iso)4e* homoeologues in the *ta-eif(iso)4e* T0 plant 2 (cv Prevert).
